# The 24-chain core-shell nanostructure of wood cellulose microfibrils in seed plants

**DOI:** 10.1101/2021.12.31.474620

**Authors:** Chih-Hui Chang, Wenjie Cai, Jer-Horng Lin, Shing-Jong Huang, Ying-Chung Jimmy Lin, Cheng-Si Tsao, Hwan-Ching Tai

**Author notes:** Correspondence: Hwan-Ching Tai, School of Public Health, Xiamen University, Xiamen, P. R. China 361102, Cheng-Si Tsao, Institute of Nuclear Energy Research, Taoyuan, R. O. C. 325.

## Abstract

Wood cellulose microfibrils (CMFs) are the most abundant organic substance on earth, but their nanostructures are poorly understood. There are controversies regarding the glucan chain number (*N*) of CMFs during initial synthesis and whether they become fused afterwards. Here, we combined small-angle X-ray scattering (SAXS), solid-state nuclear magnetic resonance (ssNMR) and X-ray diffraction (XRD) analyses to resolve these controversies. We successfully developed SAXS measurement methods for the cross-section aspect ratio and area of the crystalline-ordered CMF core, which showed higher density than the semi-disordered shell. The 1:1 aspect ratio suggested that CMFs remain mostly segregated, not fused. The area measurement revealed the chain number in the core zone (*N*_*core*_). The ratio of ordered cellulose over total cellulose, termed *R*_*oc*_, was determined by ssNMR. Using the formula *N* = *N*_*core*_ / *R*_*oc*_, we found that the majority of wood CMFs contain 24 chains, conserved between gymnosperm and angiosperm trees. The average wood CMF has a crystalline-ordered core of ∼2.2 nm diameter and a semi-disordered shell of ∼0.5 nm thickness. In naturally and artificially aged wood, we only observed CMF aggregation (contact without crystalline continuity) but not fusion (forming conjoined crystalline unit). This further argued against the existence of partially fused CMFs in new wood, overturning the recently proposed 18-chain fusion hypothesis. Our findings are important for advancing wood structural knowledge and more efficient utilization of wood resources in sustainable bio-economies.

## Introduction

Wood has been widely utilized throughout human history as fuels and building materials, also useful for making papers, utensils, and furniture ^1,2^. Nearly 300 gigatons of carbon are stored in wood, the secondary xylem of tree stems, accounting for over 50% of global biomass ^3,4^. Wood cellulose, which accounts for ∼45% of cell wall dry weight ^5^, is the most abundant organic substance on earth. As a major renewable resource in sustainable bio-economies, wood cellulose may serve as feedstock for biofuels ^6,7^ and fiber sources for novel composite materials ^8,9^. Wood with improved properties may be obtained by genetic engineering of trees ^10,11^. For these advanced applications, it is important to understand the material properties of wood cellulose, which critically depends on the organization of β-1,4-glucan chains into cellulose microfibrils (CMFs).

The nanostructures of wood CMFs remain poorly understood due to the lack of accurate methods to determine their size and shape ^12-14^. Wood CMFs are paracrystals ^15^, with crystalline-ordered chains in the center and semi-disordered chains in the periphery ^16^. The crystalline region of wood cellulose is similar to that of cellulose I_α_ or I_β_ (**Fig. 1a-1b**), so we assume that each chain occupies ∼0.320 nm^2^ of cross-section area ^17-19^. Although crystallite widths may be estimated from X-ray diffraction (XRD) data, which are around 3 nm for spruce wood ^16,20,21^, it is difficult to assess *N* because the Scherrer equation become less accurate when the crystal is semi-disordered or smaller than 4 nm ^22-24^. Visualization of wood CMFs by transmission electron microscopy (TEM) is also challenging, and the reported diameter for spruce is 2.5 nm, corresponding to just 15 chains ^25^.

**Figure 1.**
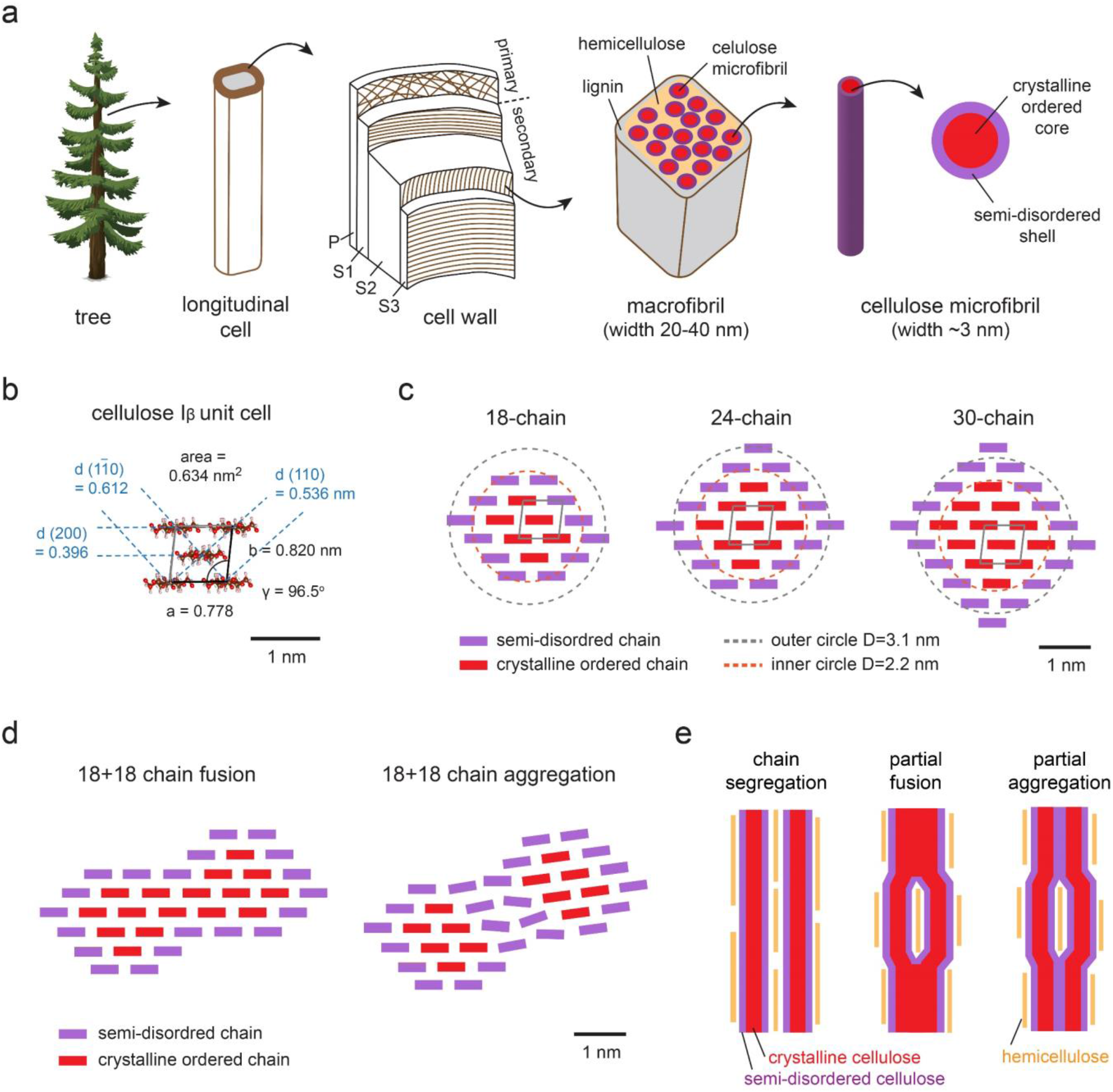
Proposed CMF structural models. (a) The great majority of wood CMFs comes from the S2 layer of longitudinal cells, with a core-shell structure. (b) Monoclinic unit cell of cellulose I_β_ ^17^. (c) CMF models with 18, 24, and 30 chains, each showing just one of many possible configurations. (d) Horizontal cross-section of a CMF pair in fusion and aggregation states. (e) Vertical cross-section of a CMF pair in segregation, partial fusion, and partial aggregation states.

It remains controversial whether CMFs stay segregated after initial synthesis or undergo coalescence, which includes fusion (forming a single crystal unit) and aggregation (lateral contact without crystalline continuity). The latest hypothesis assumes the partial fusion scenario ^26-28^ (**Fig. 1c-1e**). The second controversy concerns the number of glucan chains (*N*) per CMF during initial synthesis. According to the segregation model, the proposed values range from 18 ^29^ or 24 ^16^ for gymnosperms to 30 or 36 for angiosperms ^30^. These estimates are multiples of six because plant cellulose synthase complexes (CSCs) exhibit six-fold rosette symmetry ^31-33^. According to the fusion model, CMFs are synthesized with 18 chains but undergo fusion to increase apparent sizes, reaching *N* ∼ 22 for gymnosperms and *N* ∼ 27 for angiosperms ^26^. We shall refer to this as the 18-chain fusion model and show that it is invalid.

Instead of characterizing CMF shape and size, we achieved the breakthrough of measuring the cross-section aspect ratio and chain number (*N*_*core*_) of the crystalline-ordered core zone using small-angle X-ray scattering (SAXS). Based on the aspect ratio and XRD data, we found no evidence of CMF fusion in new or aged wood samples. Instead, CMFs remained segregated in new wood but became aggregated in millennium-old wood. We also determined the ratio of ordered cellulose over total cellulose, termed *R*_*oc*_, by solid-state nuclear magnetic resonance (NMR). Using the formula *N* = *N*_*core*_ / *R*_*oc*_, we found *N* = 24 for the majority of CMFs in both gymnosperm and angiosperm wood.

## Results and Discussions

### The geometry of the CMF core

To provide a comprehensive perspective on wood secondary cell walls, we investigated four evolutionary clades from seed plants ^34,35^. For woody gymnosperms, spruce and Chinese fir represented two common conifer families—*Pinaceae* and *Cupressaceae*, respectively. For woody angiosperms, maple and catalpa represented two major classes of eudicots—the rosids and the asterids, respectively (**SI Table S1**).

Previous studies have misconceived the use of wood SAXS patterns to model the whole CMF, and the chosen model of infinitely long cylinders was also oversimplified. This resulted in CMF diameter estimates that were too small to fit 18 chains ^25,36-38^. We showed by Porod analysis that SAXS patterns may be used to model the CMF core zone, not the whole CMF (**SI Fig. S1a**). The power-law scattering behavior with an exponent value of -4 in the high-Q region suggested the existence of a smooth interface between crystalline-ordered glucans (high-density core) and semi-disordered glucans (low-density shell). On the other hand, CMF shell glucans and hemicellulose chains only make sporadic contacts ^39,40^, unlikely to result in a smooth interface. Moreover, using simulated data, we showed that ignoring length contributions would cause an overestimation of cross-section areas (**SI Fig. S1b**, see SI for details).

Taking length contributions and potential coalescence into consideration, we modeled CMF cores using circular cylinders (CYL), elliptical cylinders (ELL), or rectangular parallelepipeds (PARA) with finite length parameters. The CYL model allowed for radius polydispersity, while ELL and PARA models could simultaneously assess the cross-section aspect ratio and area ^41^. The SAXS fitting curves for four wood species are shown in **Fig. 2a-2h** and **SI Fig. S2**, and the results are summarized in **SI Table S2**.

**Figure 2.**
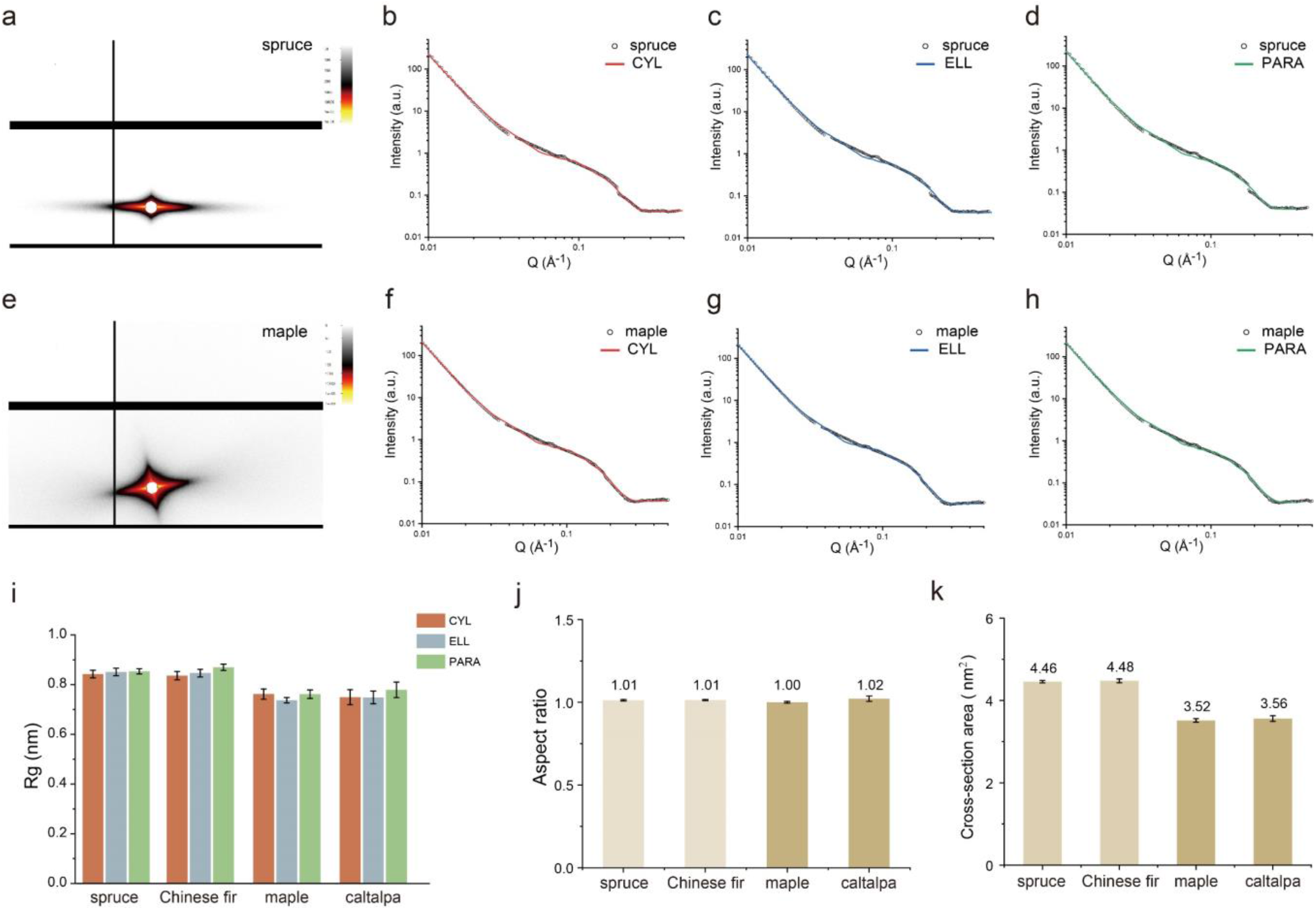
SAXS patterns of spruce (a) and SAXS profiles for comparison with fitted intensities (solid curves) using CYL (b) ELL (c) and PARA models (d). The same for maple, (e-h). Radius of gyration (R_g_) (i), aspect ratio (j), and cross-section area (k) are plotted for four wood species. Error bars represent standard error of mean (n = 9).

From the CYL fitting model, the average diameter of four wood species was 2.26 nm. For spruce, all three fitting models showed mutual agreements as they yielded similar radii of gyration; ditto for maple, fir, and catalpa (**Fig. 2i**). For all wood species, we did not observe elongated aspect ratios associated with fusion but only 1:1 aspect ratio and very low radius polydispersity, suggesting mostly segregated CMFs (**Fig. 2j**). The mean cross-section of the crystalline core, averaged over three models, was 4.46 nm^2^ for spruce (*N*_*core*_ = 13.9 chains), 4.48 nm^2^ for fir (14.0 chains), 3.52 nm^2^ for maple (11.0 chains), and 3.56 nm^2^ for catalpa (11.1 chains) (**Fig. 2k**).

### Calculating CMF chain number

The core-shell density contrast revealed by Porod analysis likely originated from differences in glucan conformational flexibility. Tightly packed glucans with crystalline order have the exocyclic C6 group locked into *tg* conformation (δC6 ∼65 ppm; δC4 ∼89 ppm) ^42,43^. By contrast, semi-disordered glucans have C6 dangling between *gt* and *gg* conformations (δC6 ∼62 ppm; δC4 ∼84 ppm), as we recently demonstrated by ^1^H-^13^C correlation spectra under ultrafast-magic angle spinning ^44^. Therefore, *N*_*core*_ / *N* should correspond to *R*_*oc*_ from NMR measurements, which is the area ratio of ordered-C4^(89 ppm)^ / total-C4^(84+89 ppm) 43,45,46^. We measured *R*_*oc*_ by spin-locking NMR experiments, which isolated cellulose subspectra based on their lower mobility (**Fig. 3, SI Fig. S3, and Table 1**).

**Table 1.** The number of glucan chains in the CMF of normal wood

| Wood species | Tree classification | Core chain number ($N_{core}$)<br>(n = 9) | Measured $R_{oc}$ (%)<br>(n = 4) | CMF chain number ($N$) |
| --- | --- | --- | --- | --- |
| spruce | gymnosperm | $13.9 \pm 0.1$ | $56.1 \pm 2.2$ | $24.8 \pm 1.0$ |
| Chinese fir | gymnosperm | $14.0 \pm 0.2$ | $55.1 \pm 1.1$ | $25.4 \pm 0.6$ |
| maple | angiosperm | $11.0 \pm 0.1$ | $46.9 \pm 0.4$ | $23.5 \pm 0.3$ |
| catalpa | angiosperm | $11.1 \pm 0.2$ | $44.5 \pm 1.4$ | $24.9 \pm 0.9$ |

Note: $R_{oc}$ is the ratio of ordered cellulose over total cellulose; $N = N_{core} / R_{oc}$ ; Values are presented as mean ± standard error; n is sample size.

**Figure 3.**
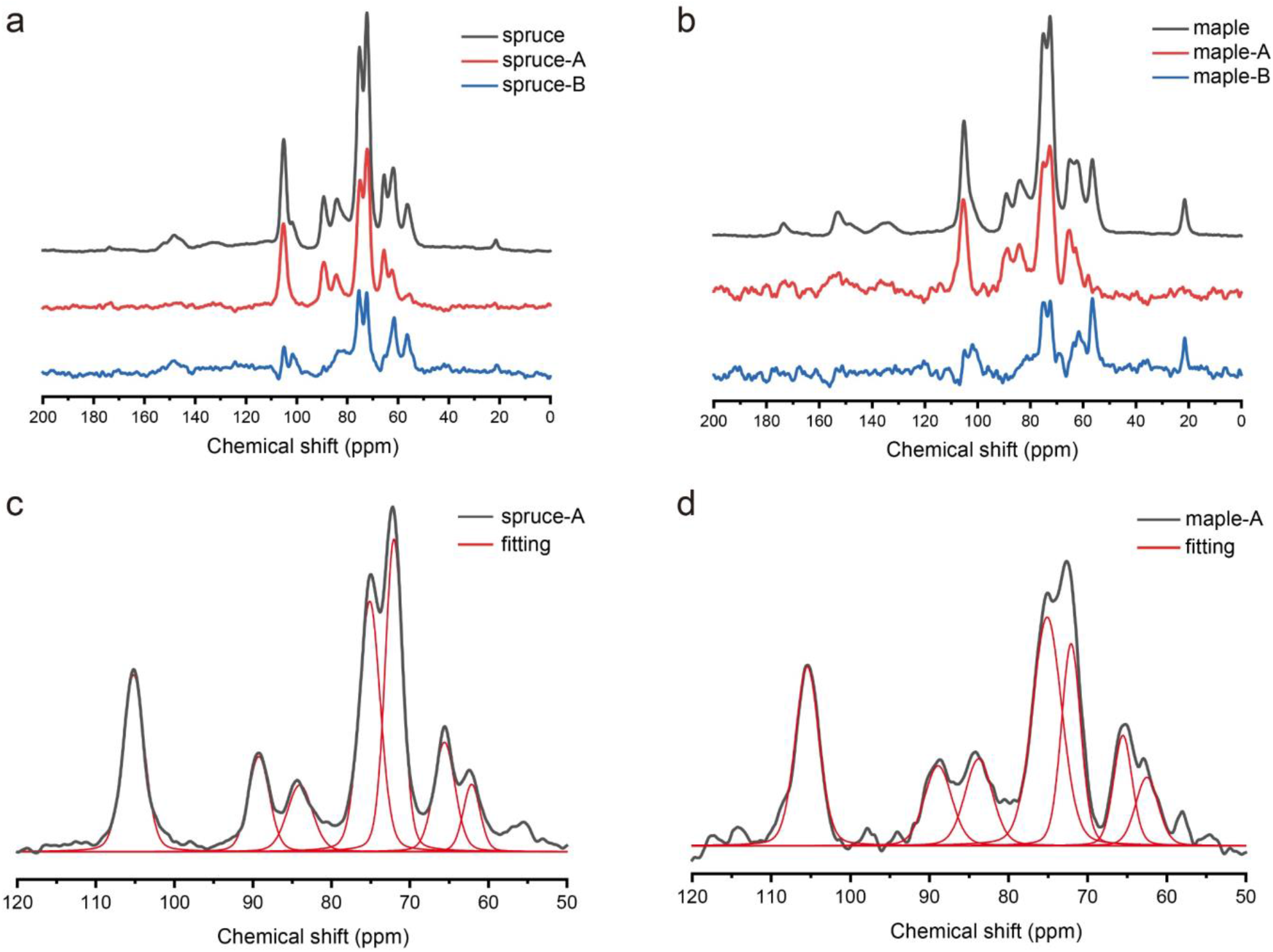
^1^H-^13^C cross-polarization spectrum of spruce (a) and maple (b), separated into subspectrum A for cellulosic components and subspectrum B for non-cellulosic components. The deconvolution of subspectrum A are shown for spruce (c) and maple (d).

Based on *N*_*core*_ values determined by SAXS, the predicted *R*_*oc*_ according to 18/24/30 chain models were 77/ 58/46% for spruce and 61/46/37% for maple, respectively. The measured *R*_*oc*_ was 56.1 ± 2.2% (mean ± standard error) for spruce and 46.9 ± 0.4% for maple, in close agreement with the 24-chain model. Using the formula *N* = *N*_*core*_ / *R*_*oc*_, the calculated CMF chain number was 24.8 ± 1.0 for spruce and 23.5 ± 0.3 for maple. The results for Chinese fir (25.4 ± 0.6) and catalpa (24.9 ± 0.9) similarly supported that the majority of CMFs in normal wood contain 24 glucan chains, conserved between gymnosperm and angiosperm trees (**Table 1**). The higher *R*_*oc*_ values of two gymnosperms over two angiosperms implied that *R*_*oc*_ is not merely determined by internal-to-surface glucan ratio but also related to the orderliness of cellulose packing, which is greater for spruce than maple ^44^.

The molecular architecture of CSCs responsible for making mature wood remains to be determined, but our data strongly suggest the involvement of 24-subunit CSCs. Our results do not rule out the possibility that some CSCs in secondary xylems may have heterogeneous compositions (18 or 30 subunits) or that some subunits may be intermittently inactive during CMF synthesis. In fact, 18-subunit CSCs ^32,47,48^ and 18-chain CMFs ^27,49^ have been suggested for non-wood plant cells. We propose that producing 24-chain CMFs in wood secondary cell walls may help provide extra mechanical support.

### Comparisons with previous reports

Spruce is the most commonly investigated wood species for CMF nanostructure, so it is useful to compare our results to literature values. Assuming a circular cylinder, our 24-chain spruce model has a core diameter of ∼2.4 nm and a shell thickness of ∼0.4 nm. We measured *R*_*oc*_ by NMR to be 56.1%, comparable to the average *R*_*oc*_ value of 56.0% from three NMR studies ^42,43,46^ and the average crystallinity index of 57.0% from four XRD studies ^50-53^. The combined width of core and shell (∼3.2 nm) was comparable to crystallite widths derived from our XRD data—3.0/3.0/3.0 nm along the directions of (110)/(1-10)/(200), respectively (**Fig. 4, SI, Fig. S4, and Table S3**). The average crystallite widths reported in the literature were 2.9/2.9/3.1 nm, respectively ^16,20,21^. Our core-shell model could explain the unstained core (2.2 nm diameter) and the semi-stained shell (0.5 nm thickness) observed in the serial TEM of de-lignified spruce ^54^. Previous SAXS studies have misreported CMF diameters to be 2.3 nm ^36^ and 2.5 nm ^25,37^, which actually corresponded to our core diameter.

**Figure 4.**
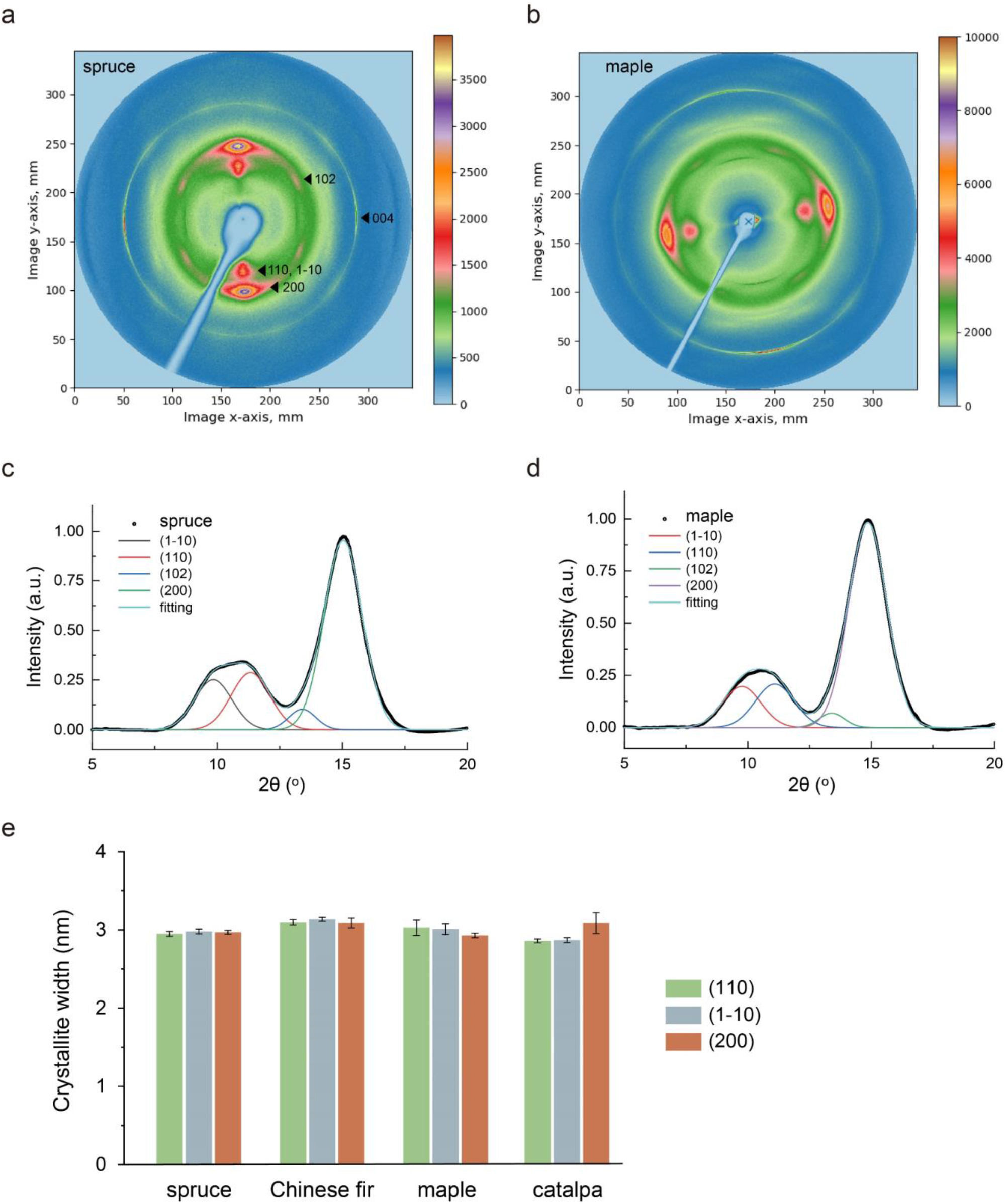
XRD patterns of spruce (a) and maple (b), with peak deconvolution analyses in (c) and (d), respectively. Crystallite widths for four wood species are plotted in (e). Error bar represents standard error of mean (n = 4).

Based on XRD and small-angle neutron scattering (SANS) data, Jarvis et al. initially proposed segregated spruce CMFs containing 24 chains ^16^ but later revised to an 18-chain fusion model with an apparent size of 22 chains ^26^. While Jarvis et al. took a heuristic approach for estimating *N*, we are the first to propose a general formula to calculate *N* that is applicable across different species, yielding 24.8 ± 1.0 (mean ± standard error) for spruce. As such, our 24-chain core-shell model provided congruent explanations for previously reported NMR, XRD, SAXS and TEM measurements. The key advantage of our analytical approach is the ability to differentiate segregation, aggregation, and fusion states, as shown below for aged wood.

### CMF aggregation in aged wood

In naturally aged wood, we hypothesized that the aspect ratio of CMF cores may increase to 2:1 due to the gradual hydrolysis of hemicellulose, allowing neighboring CMFs to coalesce (**Fig. 1e**) ^55-57^. So we investigated well-preserved Chinese fir and catalpa samples from antique Chinese guqin zithers ^58^ (**SI Table S1**). Antique firs (1200-2300 years old) exhibited an average aspect ratio of 1.8:1 and a 1.7-fold increase in cross-section area. However, there were no corresponding expansions in crystallite widths, so the coalescence was attributed to aggregation instead of fusion. By contrast, aged catalpas (∼500 years old) showed only minor signs of aggregation (**SI Fig. S5**), which may be due to younger age and/or different hemicellulose and lignin compositions between angiosperms and gymnosperms ^5^. Our data strongly suggest that CMFs in new wood are segregated, not fused. If partially fused CMFs had originally existed, it would be difficult to explain why they stopped fusing but started aggregating during aging.

In artificial aging experiments, using hot water extraction ^59^ and alkaline treatments ^57,60^ to mildly promote hemicellulose hydrolysis and/or cellulose rearrangement, we did not observe CMF fusion but only aggregation, which was more prominent in spruce than maple (**SI Fig. S6**). SANS studies have found longer center-to-center CMF distances in angiosperms (∼4.0 nm) compared to gymnosperms (3.0-3.8 nm), which were interpreted as evidence of thicker CMFs in angiosperms ^16,30,61^. However, we found similar crystallite widths between angiosperms and gymnosperms (**Fig. 4** and **Table S3**) and smaller CMF cores in the former (**Fig. 2k**). Therefore, we propose that the longer distances in angiosperms may reflect additional hemicellulose chains sandwiched between CMFs, which may explain why angiosperm CMFs are less prone to aggregation in artificial aging treatments. In sum, we found no evidence of CMF fusion in new, aged, or artificially aged wood.

## Conclusions

In this study, we demonstrated two critical misconceptions in previous studies of wood CMFs: (1) assuming post-synthesis fusion; (2) using SAXS data to model the whole CMF. By pioneering the use of SAXS to measure the aspect ratio and area of CMF cores, in combination with NMR and XRD analyses, we observed these important features: (1) *N* = 24 for the majority of wood CMFs; (2) mostly segregated CMFs in new wood; (3) similar sizes in angiosperms and gymnosperms. The average *R*_*oc*_ value of four wood species is ∼50%, which corresponds to a crystalline-ordered core with ∼2.2 nm diameter and a semi-disordered shell of ∼0.5 nm thickness. These findings overturned the recently proposed 18-chain fusion hypothesis ^26,28^.

Unfortunately, recent studies have generally accepted 18-chain CMFs as the basic building block of wood ^33,40,62,63^. It is crucial to incorporate our 24-chain core-shell model to properly explain wood nanostructures and to predict cellulose alterations due to genetic engineering or wood processing. Further research will be required to determine the cross-section chain configurations and longitudinal twists of CMFs, as well as their interactions with hemicellulose chains.

## Acknowledgement

We thank Ingo Burgert for useful manuscript discussions. Dr. Kin Woon Tong generously provided antique guqin samples, while Sandro Chiao, Boa-Tsang Lee, Dan Lu and Yu-Hsien Chu generously provided modern wood samples. We thank National Synchrotron Radiation Research Center, Taiwan for provision of beamtime at TPS-BL13A, TLS-BL23A, TLS-BL01C2 endstations. We thank NTU-AMS Laboratory for radiocarbon dating and NTU Instrument Center for NMR measurements.

