## Supplementary Information for "The 24-chain core-shell nanostructure of wood cellulose microfibrils in seed plants"

**for**

* Corresponding authors

### I. Materials and Methods

#### Wood samples

Modern and antique wood samples analyzed in this study are listed in Table S1. The modern spruce (*Picea abies*) and maple (*Acer pseudoplatanus*) came from standard European tonewood (specially selected "resonance wood") used by violin makers. The grading criteria developed by tonewood suppliers and instrument makers over the centuries allowed the selection of wood with uniform structural and mechanical properties and with greater anisotropy due to highly ordered anatomical features [1,2]. We chose tonewood-grade spruce and maple because they generally show lower sample-to-sample variations. For gymnosperm and angiosperm woods from another part of the world, we chose Chinese fir (*Cunninghamia lanceolata*) and catalpa (*Catalpa ovata*) tonewoods used by Chinese guqin (7-string zither) makers.

Antique wood samples included Chinese fir and catalpa taken from fine-sounding Chinese guqins, generously provided by Kin Woon Tong. Their species identification and radiocarbon dating results have been reported [3], with the summary shown in Table S1.

#### Artificial aging of wood

For artificial aging, wood samples were cut into thin slices about 0.5-1 mm thick, weighing around 50 mg. Hot water extraction was carried out by immersing wood slices in deionized water for 72 h at 95 ˚C. For alkaline treatments, wood slices were immersed in KOH or $Ca(OH)_2$ solution at pH 12 for 72 h at room temperature. After treatments, samples were rinsed twice with deionized water and stored in a desiccator cabinet (~40% relative humidity).

#### Synchrotron X-ray diffraction (XRD)

The XRD experiments were performed at the BL01C2 beamline of the National Synchrotron Radiation Research Center (NSRRC, Hsinchu, Taiwan), in which the ring was operated at 1.5 GeV energy with a typical current around 360 mA. A thin slice of the wood sample was fixed with 3M Magic Tape and attached to the sample holder. Two pairs of slits and one collimator were set up to provide a collimated beam with dimensions of 0.1 × 0.1 mm (H × V) at the sample. The wavelength of the incident X-rays was 1.033210 Å (12 keV), delivered from the 5-T Superconducting Wavelength Shifter and a Si (111) triangular crystal monochromator. The diffraction angles were calibrated according to Bragg positions of CeO2 (NIST SRM 674b) standards in the desired geometry.

The sample holder was placed inside a customized chamber injected with helium gas, designed to mitigate the absorption and scattering by air particles. We observed significant background signal reduction by replacing air with helium, as previously reported for protein crystallography experiments [4,5]. The incident beam passed through the wood sample in the helium gas chamber, with the beam-stop obstructing small-angle diffraction, and the other scattered light traveled toward the Mar345 image plate detector, which converted the signals into 2D pattern images after 60 s exposure time. GSAS II software (Argonne National Laboratory) was used to obtain corresponding one-dimensional powder diffraction profile with cake-type integration.

##### XRD data analysis

The XRD profile was imported into PeakFit version 4.0 (Systat Software Inc., Richmond, CA) for smoothing, background subtraction, and peak fitting. The peaks were labeled in this study according to the cellulose $I_\beta$ unit cell [6]. The background was assumed to be a quadratic function [7,8] over the 2θ range of 5°-20°. The profile over this range was fitted with four peaks, (1-10), (110), (102), and (200), following previous studies [8-10]. The peak type was Gaussian with varying widths, and adjusted until $R^2 \geq 0.995$, iteration = 7, or F-Stat ≥ 10,000.

The size of the crystalline domain (L) was estimated using the Scherrer equation: $L = K\lambda = B \cos\theta$, where λ is the X-ray wavelength, B is the full width at half maximum in 2θ units, θ is the Bragg angle, and K is the shape factor. The value of K is not known with any certainty for wood CMFs and 0.90 was chosen as an approximation, in accordance with previous studies [11-13].

##### Nuclear magnetic resonance (NMR) spectroscopy

$^{1}H$-$^{13}C$ cross-polarization (CP) NMR experiments were conducted on a wide-bore 14.1-T Bruker Avance III spectrometer equipped with a 4-mm double-resonance magic-angle spinning (MAS) probe head. The Larmor frequencies for $^{1}H$ and $^{13}C$were 600.21 MHz and 150.94 MHz, respectively. The spinning frequencies were controlled at 14 kHz. NMR rotors were filled with finely divided wood particles. For small wood sample blocks, we used small handsaws to generate sawdust. For thin-slice wood samples, we used scalpels to repeatedly cut them into very small pieces.

To obtain cellulose subspectrum from wood, we followed the spin-locking NMR experiments developed by Newman and coworkers [14,15]. Wood powders were immersed in deionized water for 7-14 days before the experiment. The delayed-contact pulse sequence consisted of a preparation time ($t_p$ = 5 μs), a spin-locking pulse ($t_{sl}$ = 6 ms), a contact time ($t_c$ = 1 ms) for CP to occur, a data acquisition time ($t_a$ = 20 ms), and a recovery time ($t_d$ = 2s) that was added before the following cycle. This sequence differed from the standard 1 ms CP-MAS sequence only by the addition of $t_{sl}$. Each sample required ~4 h of data acquisition time.

##### NMR data analysis

The subspectra of cellulose and non-cellulose components may be separated based on different mobilities, using the proton spin relaxation editing (PSRE) method [14-16]. The normal CP spectrum S was assumed to be a combination of two components, $S = A + B$, where $A$ was the cellulose subspectrum and $B$ was the non-cellulose subspectrum. Another spectrum $S'$ with spin-locking time, $t_{sl}$ = 6 ms, contained a different ratio of the two components, $S' = aA + bB$, where a and b were signal suppression factors reflecting the consequence of spin relaxation. These equations may be solved:

$$A = mS + nS'$$

$$B = (1 - m)\, S - nS'$$

, where $m = b/(b-a)$ and $n = -1/(b-a)$. The values of $a$ and $b$ followed exponential equations determined by $a = \mathrm{Exp}(-t_{sl}/T_{1\rho,A})$ and $b = \mathrm{Exp}(-t_{sl}/T_{1\rho,B})$. We adjusted $T_{1\rho,A}$ and $T_{1\rho,B}$ until the 56 ppm signal (lignin methoxy group) in the cellulose subspectrum is minimized.

The cellulose subspectrum was deconvoluted using dmfit 2020 software [17], assuming the peak shape of mixed Gaussian (80%) and Lorentzian (20%) functions. The peaks being fitted were: 61.6 ppm (C6 of non-crystalline glucans), 64.7 ppm (crystalline C6), 72.2 ppm (C2, C5), 74.8 ppm (C3, C5), 83.8 ppm (non-crystalline C4), 88.7 ppm (crystalline C4) and 104.8 ppm (C1) [18,19]. The ratio of ordered cellulose over total cellulose, termed $R_{oc}$, is defined as crystalline C4 / (non-crystalline + crystalline C4 ) = area under 89 ppm / (area under 84 ppm + area under 89 ppm), which is sometimes called the NMR crystallinity index [20,21].

**Small-angle x-ray scattering (SAXS)**

SAXS experiments were conducted at TPS-BL13A and TLS-BL23A endstations in NSRRC, Taiwan. The beam energy was 15 keV at TPS-BL13A and 8-10 keV at TLS-BL13A. The wood samples were placed in a stainless-steel holder, with exposure time of 0.1-20 s depending on sample thickness. The multi-panel detectors collected the signals of the scattered beam. SAXS data were calibrated using a custom computer program developed by NSRRC for data reduction. The SAXS profiles were modeled using the NIST analysis macros (package 7.10) [22] running under Igor Pro 6.10 (WaveMetrics, Lake Oswego, OR). The power law coefficient was set to 4. The fitting models for the wood CMFs were cylinders (CYL), elliptical cylinders (ELL), and rectangular parallelepipeds (PARA) of finite lengths. These models were developed by NIST and described in detail in the SANS model function documentation [23].

Previously, the SAXS intensity profile of a CMF was extensively modeled by the form factor of a cylinder with a radius $R$ and the assumption of infinite length [12,24-28]. The simplified model only considers the scattering contribution from the cross-section of cylinder as given by:

$$I(q) = I_0 \left( \frac{2J_1(qR)}{qR} \right)^2 \qquad \ldots\ldots\ldots\ldots..(1)$$

, where $J_1$ is the Bessel function of the first kind and first order. The scattering vector, $q$, is determined by $q = 4\pi/\lambda \sin(\theta/2)$, where $\theta$ is the SAS scattering angle and $\lambda$ is the wavelength of the incident radiation.

Because of the various orientations of CMFs (helical, orthogonal and spiral orientations), they may be randomly oriented with respect to the incident beam. Moreover, the narrow width of streak-like intensities of two-dimensional SAXS patterns demonstrated the evidence of intensity contribution from the cylinder length. Assuming that the long cylinder has the length of $L$ and the orientation of an angle $\alpha$ between cylinder axis and scattering vector, the SAXS intensity can be modelled by the form factor as given by:

$$I_{CMF}(q) = I_0 \int_0^{\frac{\pi}{2}} \left[\frac{\sin(qL/2\cos\alpha)}{(qL/2\cos\alpha)}\right]^2 \left[\frac{2J_1(qR\sin\alpha)}{(qR\sin\alpha)}\right]^2 \sin\alpha d\alpha \quad \ldots\ldots(2)$$

The usual Eq (1) only considers the scattering contribution from the cross-section of CMF. In contrast, Eq (2) considers the contribution from radius, orientation, and length. We performed a simulation work to evaluate how the neglect of length may influence the SAXS modeling result of CMFs, which has never been done for wood materials.

Figure S1b shows the calculated SAXS intensity profiles by Eq (1) for the cases of $R$ =14 and 18 Å. The Guinier shoulder of the form factor profile shifts to low $q$ values from 0.12 to 0.09 $Å^{-1}$ by increasing the $R$ value from 14 to 18 Å. The shoulder position is inversely proportional to the cylinder diameter or size. The calculated SAXS intensity profiles by Eq (2) for the cases with $L$ = 50, 500 and 10000 Å based on the same $R$ =14 Å are also shown in Figure S1b for comparison. The calculated SAXS intensity profile calculated by Eq (2) for the case with $L$ = 50 Å and $R$ =14 Å is in agreement with the $R$ = 18 Å case by Eq (1) over the main q range of 0.01 to 0.15 $Å^{-1}$. For the cases with $L$ = 500 up to 10000 Å, there is a drastic upturn, a power-law scattering behavior, in the low-q region intensity of 0.01~ 0.07 $Å^{-1}$. This power-law behavior ($I(q) \propto q^{-1}$) is mainly contributed by the length. This simulation work revealed two critical findings: (a) For the special case of CMFs with the length less than 50 Å, the approximation of Eq (1) is adequate, but this is unrealistic for wood. (b) Based on the regular cases generated by assuming $R$ = 14 Å and $L \geq 500$ Å, the model fitting result for the SAXS profile ($q \geq 0.08$ $Å^{-1}$) using Eq (1) could yield $R$ = ~18 Å, causing a significant overestimation of radius and area.

Because wood CMFs have been proposed to be partially fused [29], resulting in variable cross-section areas, we adopted the CYL model considering the circular cylinder with a finite length and polydisperse radii with Schulz size distribution *f(R)*. The SAXS intensity was calculated based on the integration of Eq (2) over radius polydispersity. Therefore, the measured SAXS profile can be modelled by

$$I(q) = Aq^{-4} + \int f(R)\ \ I_{CMF}(q)dR + b \quad \ldots\ldots\ldots\ldots\ldots\ldots\ldots\ldots\ldots\ldots(3)$$

where $A$ is a constant. The constant $b$ value is the incoherent scattering background. The first term is used to model the intensity contribution from the surface of large mesopores formed between CMFs. The upturn intensity in the measured low-$q$ region of 0.01~ 0.04 $Å^{-1}$ is mainly contributed by pore surface and has a characteristics of power-law scattering behavior ($I(q) \propto q^{-4}$). The exponent value of -4 represents the smooth pore surface according to the well-known Porod law [30]. The measured SAXS profiles were well-fitted by Eq (3).

If there were partial fusion or aggregation, CMF cross-sections may become elongated. Therefore, we adopted the ELL model corresponding to a cylinder with elliptical cross-sections. It considered randomly oriented cylinders with finite length and uniform density. An alternative to an elliptical cross-section could be a rectangular one. Hence, we also adopted the PARA model corresponding to a rectangular parallelepiped.

The radius of gyration ($R_g$) of an object instead of its diameter or width is frequently used to quantitatively characterize the size regardless of their shape. One of the advantages using radius of gyration is that shape information is not required. The $R_g$ value is usually determined by the direct Guinier approximation, a model-independent approach. This work did not adopt the Guinier approximation due to the superposition of complex contributions. We adopted the radius of gyration of a cross-section calculated by the model-fitting results. The average of the calculated $R_g$ values based on different models and/or shapes can be an effective judgement of cross-section size, minimizing the errors from the assumption of possible shapes. To date, the realistic shape of CMF cross-sections or its core has not been determined. A circular cross-section is the most adopted model.

$R_g$ is defined as the root mean square distance from the center of mass:

$$R_g^2 = \frac{\int_{V_p} \Delta\rho(r) r^2 dr}{\int_{V_p} \Delta\rho(r) dr} \quad \ldots\ldots\ldots\ldots\ldots\ldots\ldots\ldots\ldots\ldots (4)$$

where $\Delta\rho(r)$ is the contrast or difference between scattering length densities (SLDs) of the investigated particle and the surrounding material (closely related to electron density), and $V_p$ is the volume of the particle.

In the case of a circular cylinder with radius $R$ and length $L$, the $R_g$ corresponding to 3D cylinder is described by:

$$R_g^2 = \frac{R^2}{2} + \frac{L^2}{12} \quad \ldots\ldots\ldots\ldots\ldots\ldots\ldots\ldots\ldots\ldots (5)$$

The degree of polymerization of wood cellulose is ~10,000 [31], which corresponds to CMF length of ~5 μm. Our model-fitted lengths of 30000-70000 Å were consistent with this value. The q-range measured in this study was insufficient to provide accurate estimate of the full $R_g$ value of CMF including the length. The SAXS profile of wood showed a shoulder around q = 0.1 $Å^{-1}$, corresponding to structural features or form factor of the cross-section of CMF cores. Therefore, we only considered the $R_g$ value calculated from the contribution of the cross-section but not the length. Hence, for the circular cylinder, $R_g^2 = R^2/2$. For an elliptical cylinder with axes A and B, $R_g^2 = (A^2+B^2)/4$. For a rectangular parallelepiped with side lengths x and y, $R_g^2 = (x^2+y^2)/12$.

There were confusions in the literature on whether the cylinder diameter determined by SAXS analysis is the CMF width (core and shell combined) or just the core width [12,24-28]. The present SAXS study provided in-depth analysis for solving this confusion. Generally, the SAXS is an effective tool for the two-phase system. In order to use SAXS profile to model the CMF core, the SLD of the crystalline-ordered core has to be different from that of the semi-disordered shell, leading to the significant scattering contrast and thus SAXS intensity.

The accurate SLD values of the core zone, shell zone, and hemicellulose matrix are very difficult to estimate. We proposed the following strategy to justify the core diameter as the modeled diameter. If the SLD contrast detected by SAXS is dominated by the cellulose core/shell interface, it should correspond to a smooth surface characterized by the high-q profile of SAXS with a power-law scattering behavior of $I(q) \propto q^{-4}$. If the SLD contrast detected by SAXS is dominated by the shell zone/hemicellulose interface, it should correspond to a rough surface characterized by: $I(q) \propto q^{-\alpha}$, $3<\alpha<4$, with surface fractal dimension = $6-\alpha$. Recent solid-state NMR studies have shown that hemicellulose chains only make sporadic contacts with the CMF shell [32,33], which should result in rough interface morphology.

Porod plot ($I(q)\times q^4$ vs $q$) is usually adopted to examine whether the characteristics of smooth interface/surface exist according to the Porod law. The Porod plot of SAXS profile of maple via subtraction from an appropriate background value is shown in Fig. S1a. The curve demonstrates a plateau in the high-q range (0.17~0.25 Å), representing the power-law scattering behavior of $I(q) \propto q^{-4}$, consistent with the smooth interface at the core/shell boundary. The Porod plot provided the evidence of SAXS intensity contributed by the core diameter. Therefore, in our two-phase model, the first phase is the denser crystalline-ordered core, and the second phase consists of the semi-disordered shell and the hemicellulose matrix.

#### II. Tables

**Table S1.** Modern and aged wood samples analyzed

| Modern tonewood samples | | | | |
|---|---|---|---|---|
| Wood type | Identified species | Common name | Classification | Place of origin |
| spruce | *Picea abies* | Norway spruce | softwood | Europe |
| Chinese fir | *Cunninghamia lanceolata* | *shan* | softwood | China |
| maple | *Acer pseudoplatanus* | sycamore maple | hardwood | Europe |
| catalpa | *Catalpa ovata* | *zi* | hardwood | China |
| **Aged wood samples from antique Chinese zithers** | | | | |
| Sample No. | Identified species | $^{14}$C age (year before present) | Error (±) | Median calendar year |
| T1-1 | *Cunninghamia lanceolata* | 1352 | 61 | 683 |
| T1-4 | *Cunninghamia lanceolata* | 1213 | 40 | 816 |
| T3-2 | *Cunninghamia lanceolata* | 2242 | 75 | 279 BC |
| T1-2 | *Catalpa ovata* | 373 | 60 | 1536 |
| T4-4 | *Catalpa ovata* | 285 | 55 | 1584 |
| T5-1 | *Catalpa ovata* | 422 | 36 | 1465 |

**Table S2.** SAXS model fitting results for four wood species

| Model | Fitting result | Spruce | Chinese fir | Maple | Catalpa |
|---|---|---|---|---|---|
| CYL | radius (nm) | 1.19 | 1.18 | 1.08 | 1.06 |
| CYL | radius polydispersity | 1.05E-3 | 4.37E-4 | 7.71E-4 | 4.84E-4 |
| CYL | cross-section area ($nm^2$) | 4.46 | 4.40 | 3.65 | 3.54 |
| CYL | length (nm) | 4.03E+3 | 1.36E+3 | 4.52E+3 | 4.57E+3 |
| CYL | radius of gyration (nm) | 0.84 | 0.84 | 0.76 | 0.75 |
| ELL | long axis (nm) | 1.21 | 1.20 | 1.05 | 1.07 |
| ELL | short axis (nm) | 1.20 | 1.20 | 1.03 | 1.05 |
| ELL | aspect ratio (long/short) | 1.01 | 1.00 | 1.02 | 1.02 |
| ELL | cross-section area ($nm^2$) | 4.55 | 4.50 | 3.42 | 3.52 |
| ELL | length (nm) | 2.94E+3 | 2.28E+3 | 7.13E+3 | 7.17E+3 |
| ELL | radius of gyration (nm) | 0.85 | 0.85 | 0.74 | 0.75 |
| PARA | long side (nm) | 2.12 | 2.16 | 1.88 | 1.97 |
| PARA | short side (nm) | 2.07 | 2.10 | 1.85 | 1.85 |
| PARA | aspect ratio (long/short) | 1.03 | 1.03 | 1.02 | 1.07 |
| PARA | cross-section area ($nm^2$) | 4.37 | 4.52 | 3.48 | 3.63 |
| PARA | length (nm) | 3.43E+3 | 2.89E+3 | 6.96E+3 | 6.35E+3 |
| PARA | radius of gyration (nm) | 0.85 | 0.87 | 0.76 | 0.78 |
| CYL+ELL +PARA | mean area ($nm^2$)* | 4.46 | 4.47 | 3.52 | 3.56 |
| CYL+ELL +PARA | std. err. of mean area | 0.03 | 0.05 | 0.04 | 0.07 |

Note: The sample size is 9 for each wood species. *The area estimate for each sample is the average from three models.

**Table S3**. XRD fitting results and crystallite widths for four wood species

| Analysis type | Fitting result | Spruce | Chinese fir | Maple | Catalpa |
|---|---|---|---|---|---|
| (110) peak | 2θ (°) | 11.34 | 11.23 | 11.10 | 11.27 |
| (110) peak | FWHM (°) | 1.82 | 1.73 | 1.77 | 1.87 |
| (110) peak | d-spacing | 0.523 | 0.529 | 0.534 | 0.526 |
| (110) peak | Scherrer width (nm) | 2.95 | 3.10 | 3.03 | 2.86 |
| (110) peak | std. err. of mean width | 0.03 | 0.04 | 0.10 | 0.03 |
| (1-10) peak | 2θ (°) | 9.82 | 9.73 | 9.79 | 9.90 |
| (1-10) peak | FWHM (°) | 1.79 | 1.71 | 1.78 | 1.86 |
| (1-10) peak | d-spacing (nm) | 0.604 | 0.609 | 0.605 | 0.599 |
| (1-10) peak | Scherrer width (nm) | 2.98 | 3.14 | 3.01 | 2.87 |
| (1-10) peak | std. err. of mean width | 0.03 | 0.02 | 0.07 | 0.03 |
| (200) peak | 2θ (°) | 15.03 | 15.09 | 14.89 | 14.94 |
| (200) peak | FWHM (°) | 1.81 | 1.74 | 1.83 | 1.75 |
| (200) peak | d-spacing (nm) | 0.395 | 0.393 | 0.399 | 0.397 |
| (200) peak | Scherrer width (nm) | 2.97 | 3.09 | 2.93 | 3.09 |
| (200) peak | std. err. of mean width | 0.02 | 0.06 | 0.03 | 0.14 |
| Unit cell* | cross-section area per chain (nm) | 0.316 | 0.321 | 0.323 | 0.315 |

Note: FWHM is full width at half maximum. Sample size is 4 for each wood species; *calculated from d-spacing values mapped onto the monoclinic unit cell.

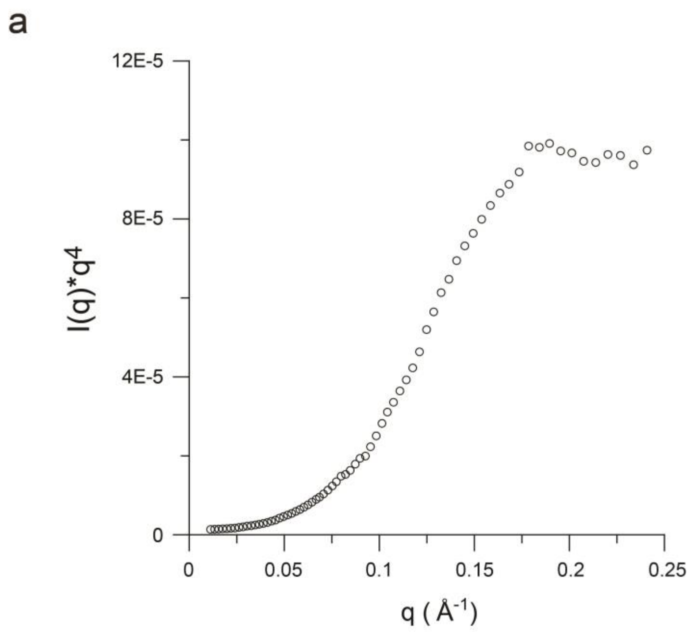


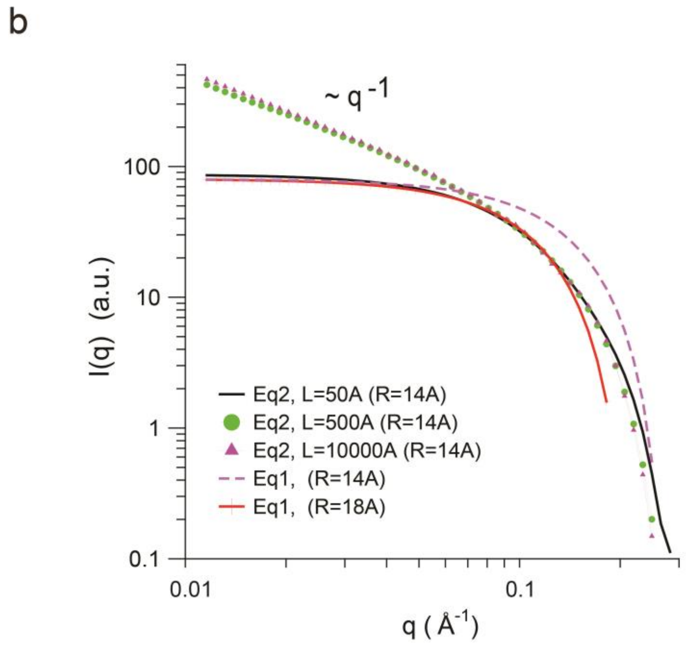


**Figure S1**. (a) Porod analysis for maple SAXS profile. (b) The SAXS profiles calculated by Eq (1) based on different radii, in comparison with those calculated by Eq (2) based on the same radius but different lengths.

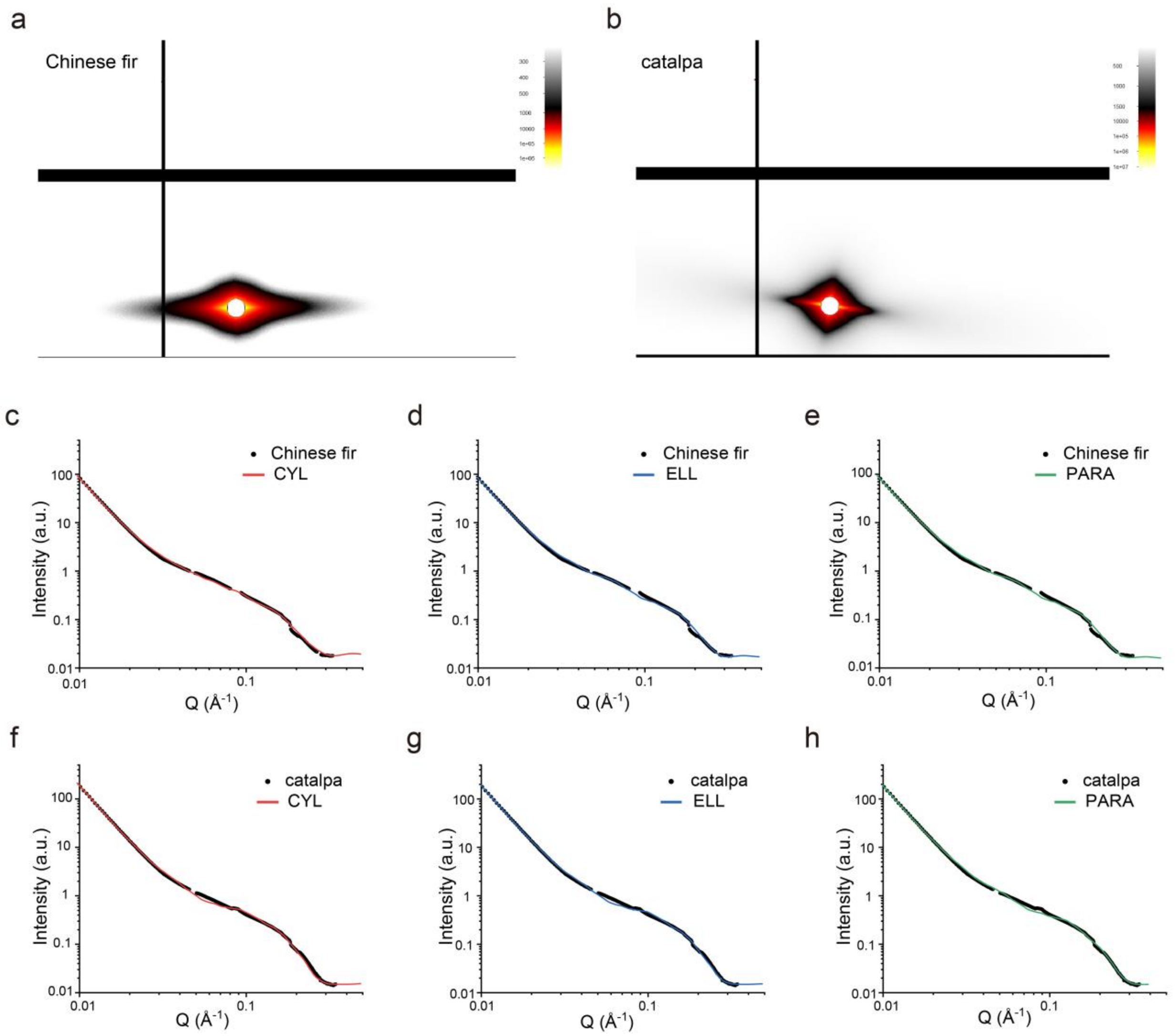


**Fig S2**. SAXS profiles of Chinese fir (a) and its curve fitting with CYL (c) ELL (d) and PARA models (e). The same are shown for catalpa (b) and its fitting results (f-h).

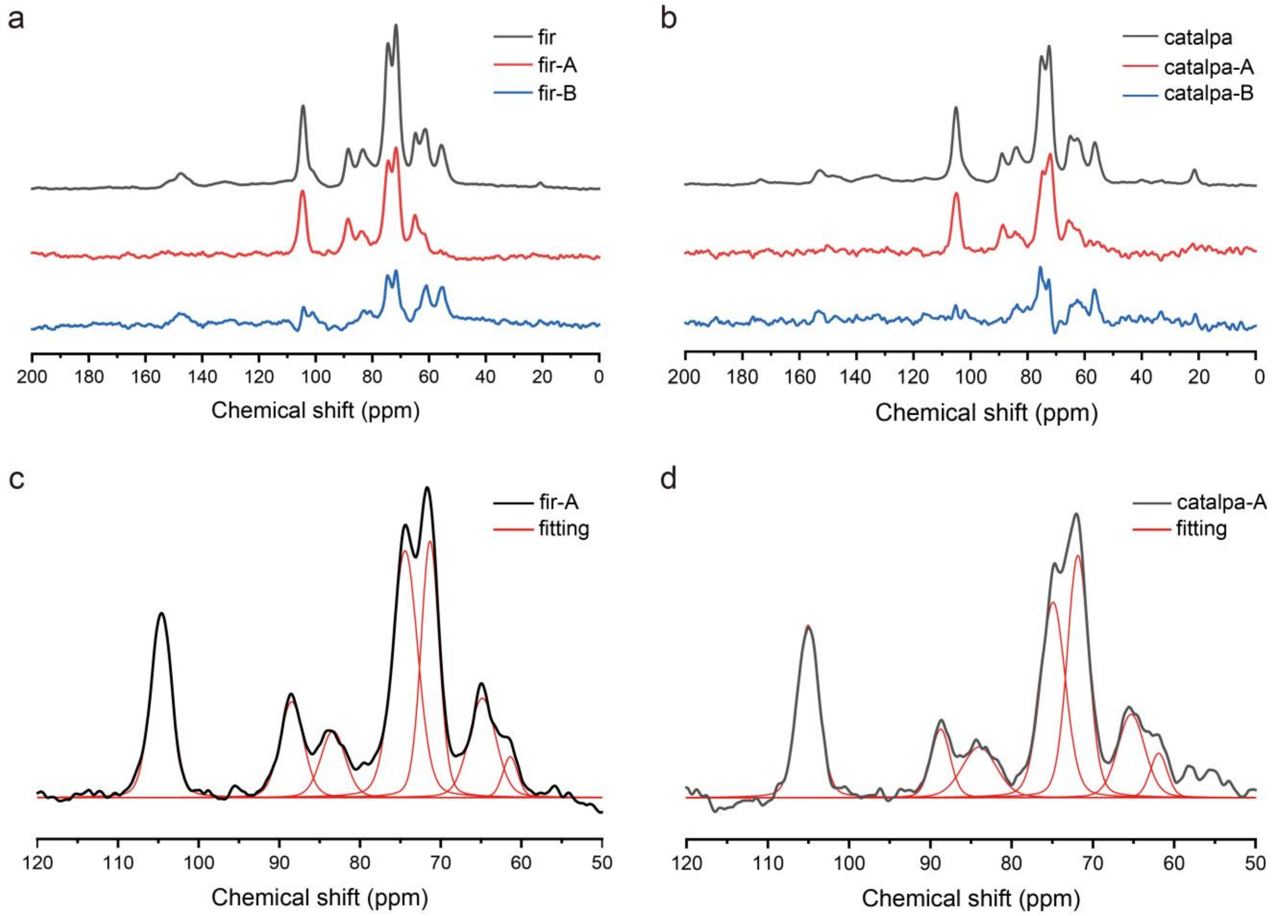


**Fig S3** $^{1}H$-$^{13}C$ cross-polarization spectrum of Chinese fir (a) and catalpa (b), separated into subspectrum A for cellulosic components and subspectrum B for non-cellulosic components. The deconvolution of subspectrum A are shown for Chinese fir (c) and catalpa (d).

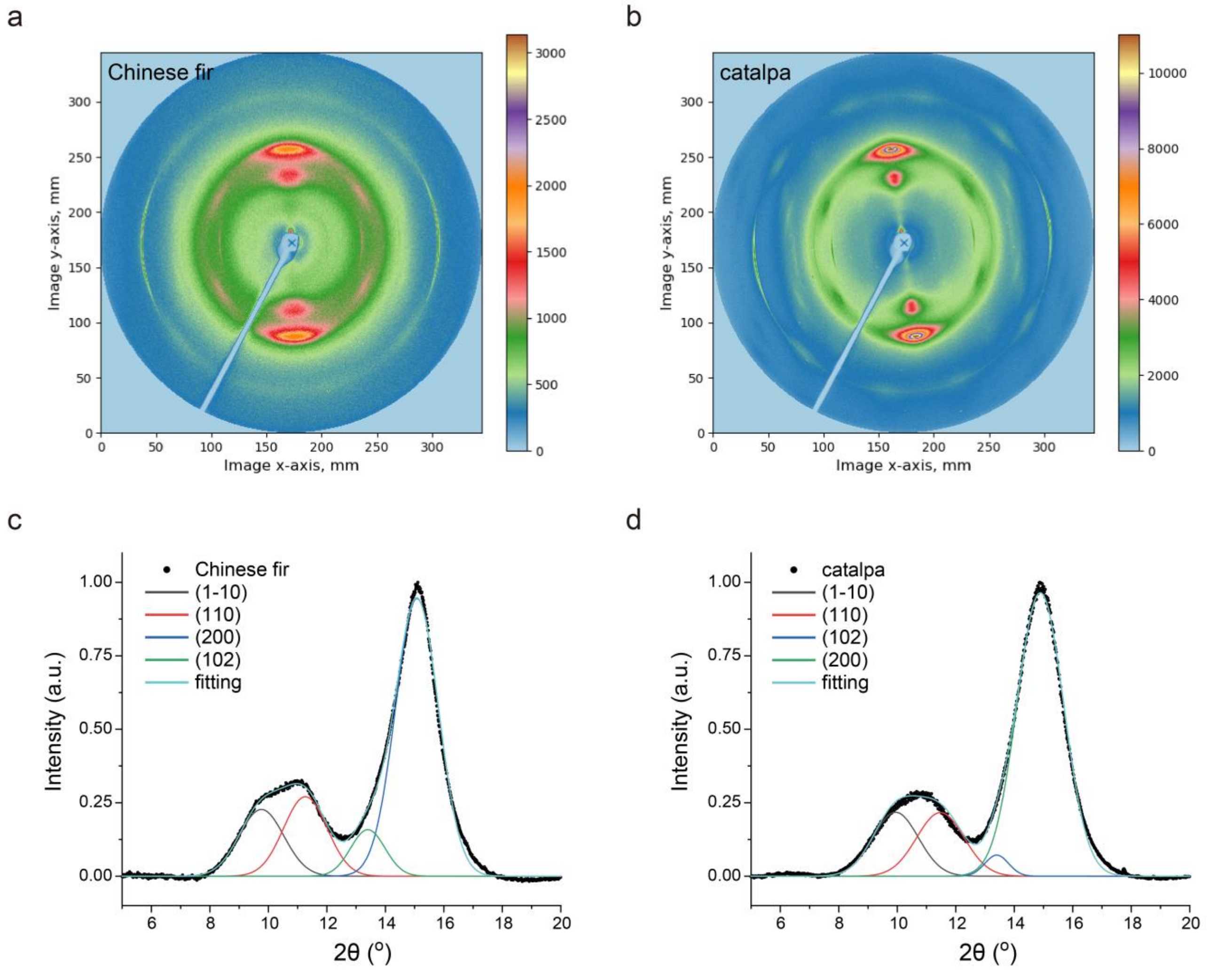


**Fig S4**. XRD patterns of Chinese fir (a) and catalpa (b), with peak deconvolution analyses in (c) and (d), respectively.

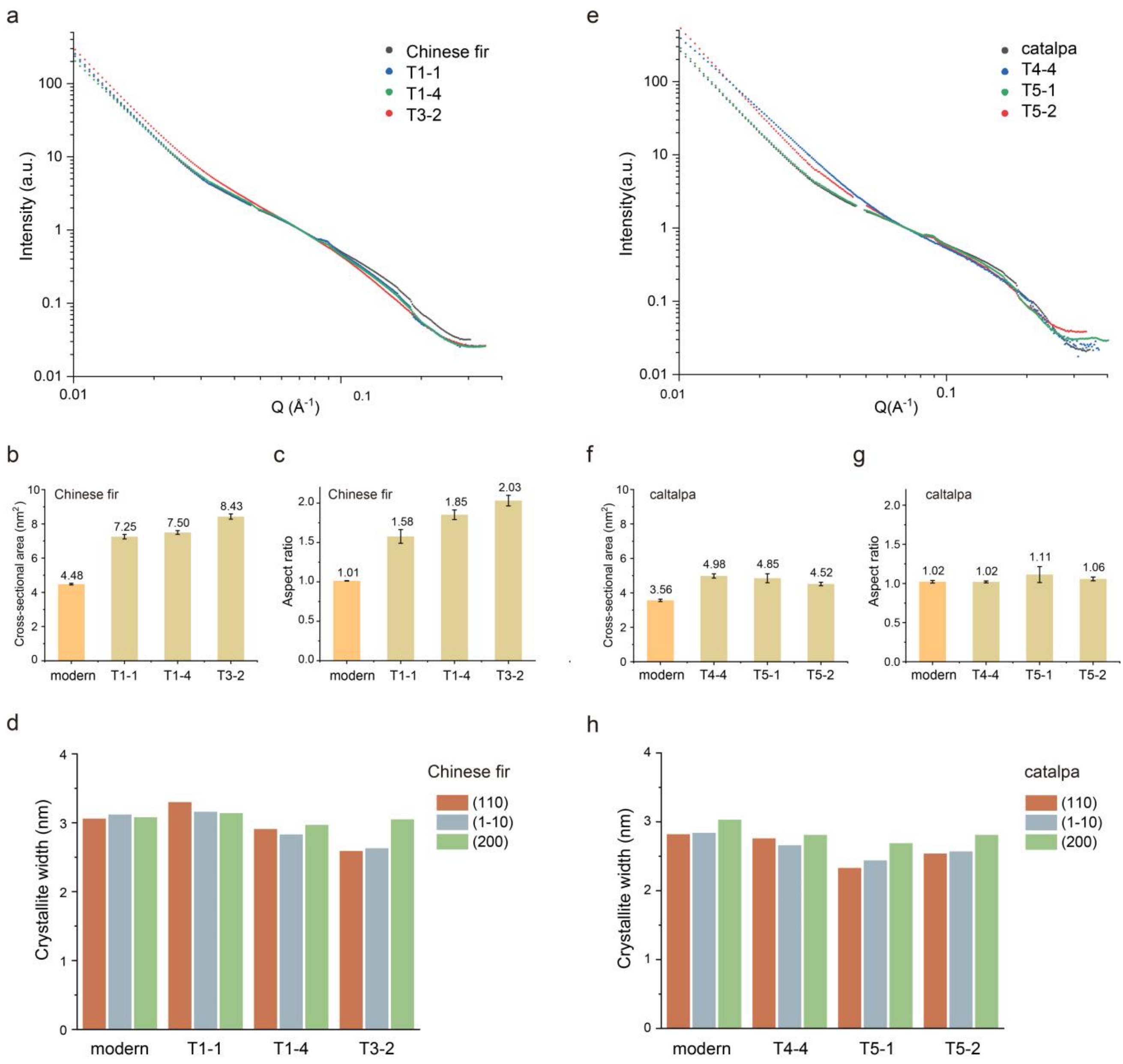


**Fig. S5**. Comparisons between modern and naturally aged woods, including Chinese fir and catalpa. SAXS profile comparisons between modern and aged firs are shown in (a). We plotted the cross-section area (b), the cross-section aspect ratio (c), and the crystallite widths along the directions of (110), (1-10), and (200) (d). Error bar represents standard error of mean (modern n=9; aged n=3). The corresponding plots for modern and aged catalpas are shown in (e-h).

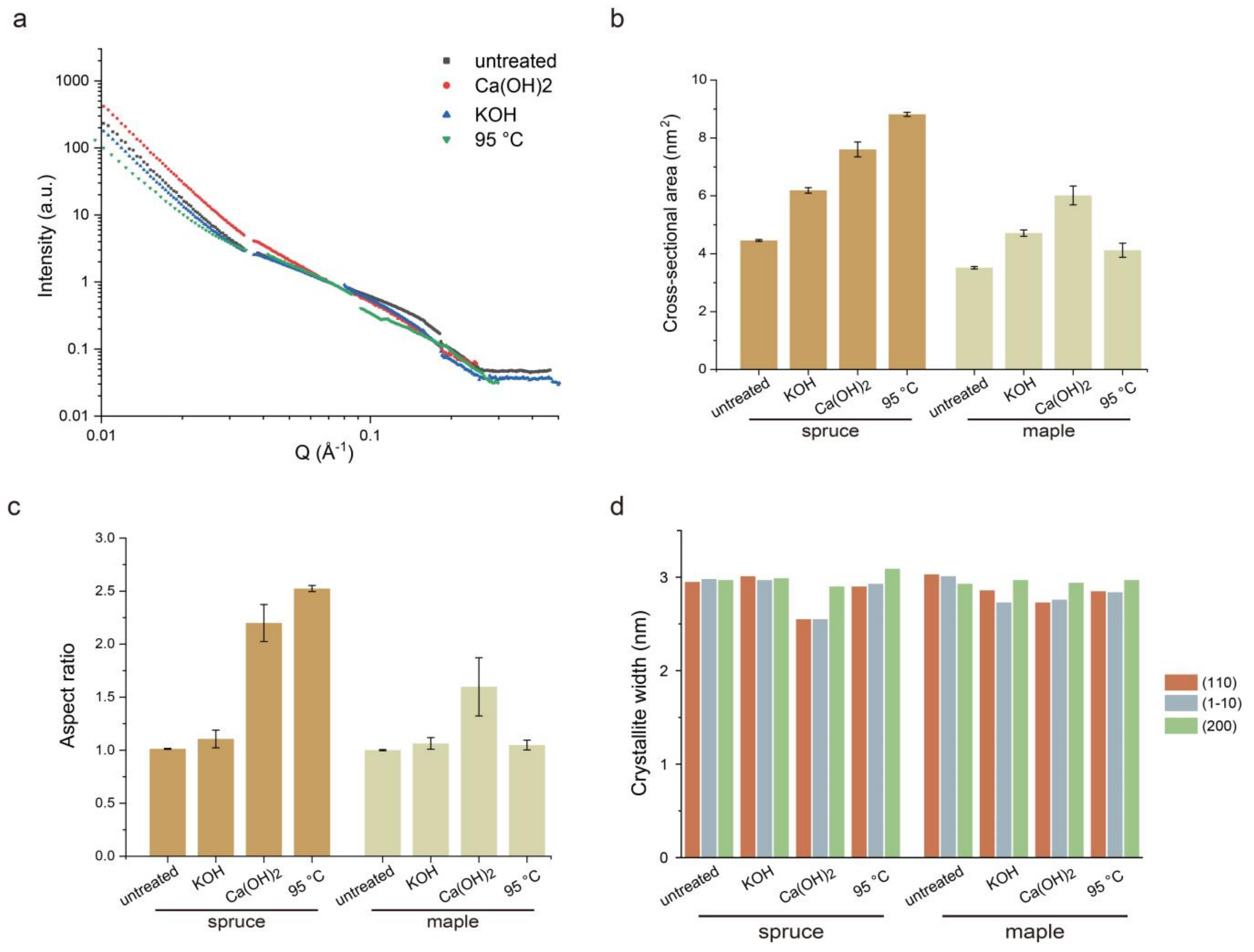


**Fig. S6**. Comparisons between modern and artificially aged woods, including spruce and maple. SAXS profile comparisons for untreated spruce and spruce treated with KOH, $Ca(OH)_2$, and 95 ˚C water are shown in (a). We plotted the area (b) and the aspect ratio (c) of the core cross-section, and the crystallite widths along the directions of (110), (1-10), and (200) (d). Error bar represents standard error of mean (untreated n=9; treated n=3).

16 Newman, R. H. & Condron, L. M. Separating subspectra from cross-polarization magic-angle spinning nuclear magnetic resonance spectra by proton spin relaxation editing. *Solid State Nucl.*

*Magn. Reson.* **4**, 259-266 (1995).
17 Massiot, D. *et al.* Modelling one-and two-dimensional solid-state NMR spectra. *Magn. Reson. Chem.* **40**, 70-76 (2002).
18 Mori, T. *et al.* Exploring the conformational space of amorphous cellulose using NMR chemical shifts. *Carbohydr. Polym.* **90**, 1197-1203 (2012).
19 Kono, H. *et al.* CP/MAS 13C NMR study of cellulose and cellulose derivatives. 1. Complete assignment of the CP/MAS 13C NMR spectrum of the native cellulose. *J. Am. Chem. Soc.* **124**, 7506-7511 (2002).
20 Teeäär, R., Serimaa, R. & Paakkarl, T. Crystallinity of cellulose, as determined by CP/MAS NMR and XRD methods. *Polym. Bull.* **17**, 231-237, doi:10.1007/BF00285355 (1987).
21 Bourmaud, A. *et al.* Evolution of flax cell wall ultrastructure and mechanical properties during the retting step. *Carbohydr. Polym.* **206**, 48-56 (2019).
22 Kline, S. R. Reduction and analysis of SANS and USANS data using IGOR Pro. *J. Appl. Crystallogr.* **39**, 895-900 (2006).
23 Kline, S. R. *SANS Model Function Documentation*. (https://github.com/sansigormacros/docs/blob/main/SANS_Model_Docs.pdf, 2012).
24 Jakob, H., Fratzl, P. & Tschegg, S. Size and arrangement of elementary cellulose fibrils in wood cells: a small-angle X-ray scattering study of Picea abies. *J. Struct. Biol.* **113**, 13-22 (1994).
25 Paris, O., Zollfrank, C. & Zickler, G. A. Decomposition and carbonisation of wood biopolymers—a microstructural study of softwood pyrolysis. *Carbon* **43**, 53-66 (2005).
26 Jungnikl, K., Paris, O., Fratzl, P. & Burgert, I. The implication of chemical extraction treatments on the cell wall nanostructure of softwood. *Cellulose* **15**, 407-418 (2008).
27 Penttilä, P. A., Rautkari, L., Österberg, M. & Schweins, R. Small-angle scattering model for efficient characterization of wood nanostructure and moisture behaviour. *J. Appl. Crystallogr.* **52**, 369-377 (2019).
28 Viljanen, M., Ahvenainen, P., Penttilä, P., Help, H. & Svedstrm, K. Ultrastructural X-ray scattering studies of tropical and temperate hardwoods used as tonewoods. *Iawa J.* **41**, 1-19 (2020).
29 Jarvis, M. C. Structure of native cellulose microfibrils, the starting point for nanocellulose manufacture. *Philos. Trans. A. Math. Phys. Eng. Sci.* **376**, 20170045, doi:10.1098/rsta.2017.0045 (2018).
30 Glatter, O. & Kratky, O. *Small-Angle X-Ray Scattering*. (Academic Press, 1981).
31 Pettersen, R. C. in *The Chemistry of Solid Wood Advances in Chemistry* Ch. 2, 57-126 (American Chemical Society, 1984).
32 Kang, X. *et al.* Lignin-polysaccharide interactions in plant secondary cell walls revealed by solid-state NMR. *Nat. Commun.* **10**, 347, doi:10.1038/s41467-018-08252-0 (2019).
33 Terrett, O. M. *et al.* Molecular architecture of softwood revealed by solid-state NMR. *Nat. Commun.* **10**, 4978, doi:10.1038/s41467-019-12979-9 (2019).
